# Structural variants in Chinese population and their impact on phenotypes, diseases and population adaptation

**DOI:** 10.1101/2021.02.09.430378

**Authors:** Zhikun Wu, Zehang Jiang, Tong Li, Chuanbo Xie, Liansheng Zhao, Jiaqi Yang, Shuai Ouyang, Yizhi Liu, Tao Li, Zhi Xie

**Affiliations:** State Key Laboratory of Ophthalmology, Zhongshan Ophthalmic Center, Sun Yat-sen University, Guangzhou, China; Sun Yat-sen University Cancer Center, Sun Yat-sen University, Guangzhou, China; Mental Health Center and Psychiatric Laboratory, the State Key Laboratory of Biotherapy, West China Hospital of Sichuan University, Chengdu, China; Guangdong-Hong Kong-Macao Greater Bay Area Center for Brain Science and Brain-Inspired Intelligence, Guangzhou, China

## Abstract

A complete characterization of genetic variation is a fundamental goal of human genome research. Long-read sequencing (LRS) improves the sensitivity for structural variant (SV) discovery and facilitates a better understanding of the SV spectrum in human genomes. Here, we conduct the first LRS-based SV analysis in Chinese population. We perform whole-genome LRS for 405 unrelated Chinese, with 68 phenotypic and clinical measurements. We discover a complex landscape of 132,312 non-redundant SVs, of which 53.3% are novel. The identified SVs are of high-quality validated by the PacBio high-fidelity sequencing and PCR experiments. The total length of SVs represents approximately 13.2% of the human reference genome. We annotate 1,929 loss-of-function SVs affecting the coding sequences of 1,681 genes. We discover new associations of SVs with phenotypes and diseases, such as rare deletions in *HBA1*/*HBA2/HBB* associated with anemia and common deletions in *GHR* associated with body height. Furthermore, we identify SV candidates related to human immunity that differentiate sub-populations of Chinese. Our study reveals the complex landscape of human SVs in unprecedented detail and provides new insights into their roles contributing to phenotypes, diseases and evolution. The genotypic and phenotypic resource is freely available to the scientific community.

## Introduction

Human genetic variants comprise single-nucleotide variants (SNVs), small insertions or deletions (InDels) and structural variants (SVs), and profoundly contribute to many physical traits and human diseases. SVs are defined as genomic rearrangements that range from 50 base-pairs (bp) to over megabases (Mbs) in length, including different forms such as deletion (DEL), insertion (INS), duplication (DUP) and inversion (INV)^1^. Accumulating evidence has demonstrated that SVs are associated with many human diseases, such as neurodevelopment disease and cancer^2–6^.

While substantial progress had been made in uncovering SNVs and InDels based on short-read sequencing (SRS) technologies, the discovery and genotyping of SVs have been hampered due to the limited power of SRS to detect SVs that frequently occur in repetitive regions with complex structures. Consequently, the current human reference genome contains a comprehensive map of SNVs, InDels, but only a relatively small number of SVs^7^. More recently, third-generation sequencing (TGS) platforms such as Pacific Biosciences (PacBio) and Oxford Nanopore Technologies (ONT) provide long-read sequencing (LRS), which improves the sensitivity for SV discovery and facilitates a better understanding of the SV spectrum in human genomes^8,9^.

Recently, a milestone study generated 15 human genomes using LRS with PacBio technology^8^. Despite the small sample size, the authors discovered 99,604 non-redundant SVs and 2,238 SVs were shared by all 15 genomes. More recently, a population-scale SV study using LRS was reported on an Icelandic population^10^. The authors identified a median of 23,111 SVs in each sample and some of them might be Icelander specific. Icelanders are a North Germanic ethnic group that is historically coherent, with an estimated population close to 360,000. In contrast, Han Chinese is the largest ethnic group in the world, with a total population of more than 1.3 billion, comprising around 19% of the human population. China has an area of 9.6 million km^2^, with two mega-regions, North and South China. Although several recent studies have reported on SVs in Chinese genomes using LRS^9,11–13^, none of them depicted multiple genomes. The complexity and diversity of genetic variation characterized by SVs among Chinese populations are unclear.

In order to gain a more complete view of human genomes, we genotyped Chinese population by performing whole-genome LRS of 405 unrelated Chinese individuals, with 68 phenotypic and clinical measurements. We detected 132,312 non-redundant SVs, of which 53.3% were novel. The identified SVs were of high-quality validated by the PacBio high fidelity (HiFi) sequencing and PCR experiments. We showed the global properties of the SVs and their widespread phenotypic, clinical and evolutionary impacts. To sum, we present an important resource for human genome research and precision medicine. Our study reveals the complex landscape of human SVs in unprecedented detail and provides new insights into their roles contributing to phenotypes, diseases and evolution. The genotypic and phenotypic resource is freely available to the scientific community.

## Results

### Sequencing, SV discovery and validation

We performed whole-genome LRS for 405 unrelated Chinese via the PromethION platform (ONT). Among all the individuals, 206 (50.9%) were males and 199 (49.1%) were females. The ages ranged from 22 to 81 years, with a median age of 42 years. These individuals were from 18 provinces in the North (124 individuals), South (198) and Southwest (53) of China, and the ancestral regions of 30 individuals were not known (**Fig. 1a** and **Supplementary Table 1**). Among them, 68 phenotypic and clinical measurements for 327 individuals were obtained by health screening (**Supplementary Table 2**).

**Figure 1.**
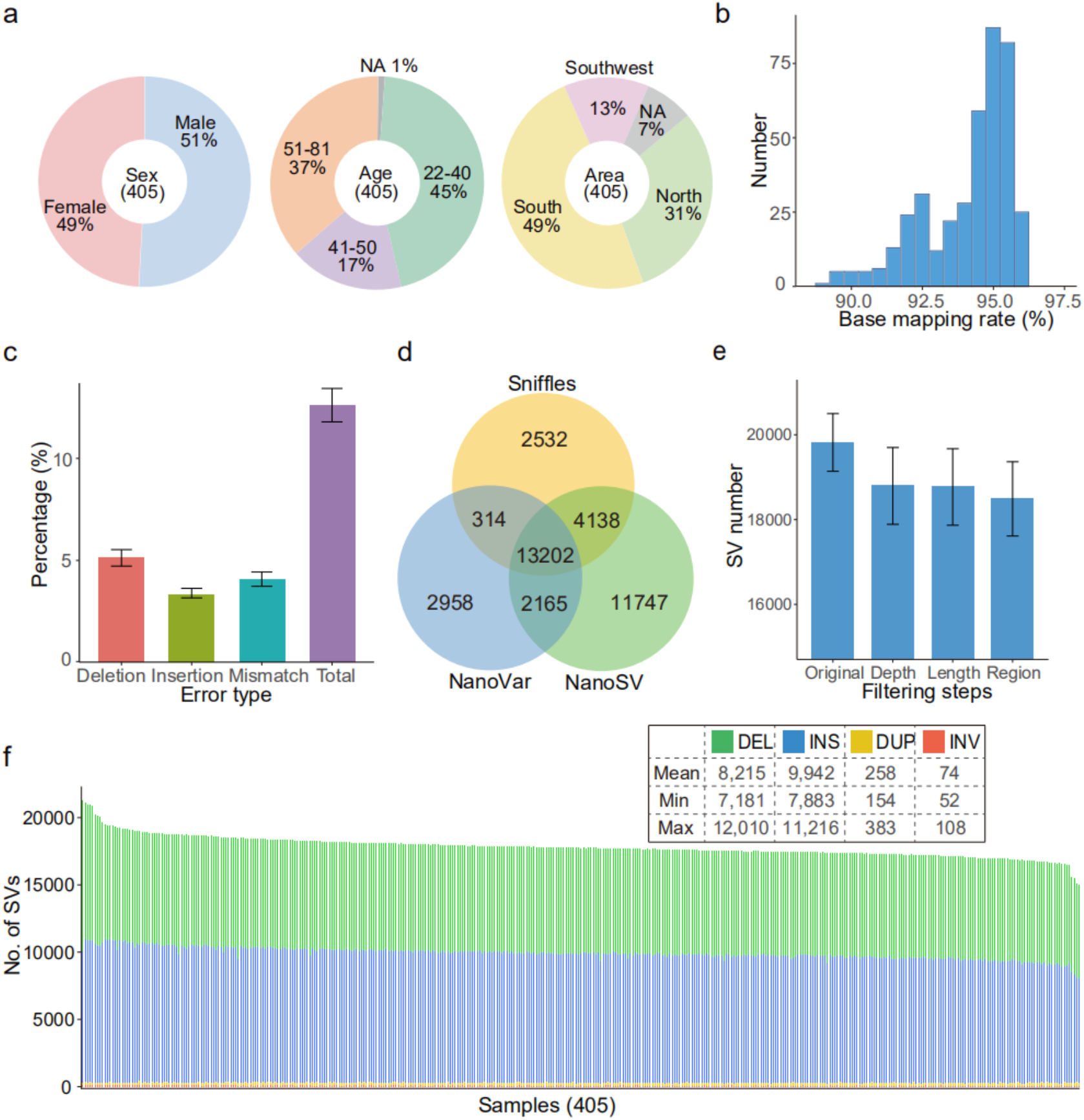
Overview of samples, datasets and SVs. **a**, Overview of information of samples in the study, “NA” denotes not available. **b**, Distribution of base mapping rate of clean data aligning to the reference genome GRCh38. **c**, Error rate for each type, error bars represent standard deviation. **d**, Average No. of SVs identified by Sniffles, NanoVar and NanoSV per individual and the overlaps among them. **e**, Average No. of SVs after each filtering step, “Original” means unfiltered SVs detected by at least two callers, “Depth”, “Length” and “Region” mean SVs after being filtered by supported reads number, extra-long interval and very low complex regions, respectively. **f**, No. of SV for each SV type in each individual.

Resequencing of these 405 Chinese generated a total of 20.7 terabases (Tb) of cleaned sequences, with an average of 51.0 gigabases (Gb) per individual, representing an average depth of approximate 17-fold (**Supplementary Fig. 1a** and **Supplementary Table 1**). The cleaned reads were then mapped onto the human reference genome GRCh38, and base mapping rate for different individuals varied from 89.0% to 96.2%, with an average of 94.1% (**Fig. 1b**). The mean error rate was 12.6%, ranging from 10.8% to 16.0% (**Methods**), which was lower than that of a recent study (15.2%)^10^ and similar to a prior benchmarking study (12.6%)^14^. The percent of deletions, insertions and substitutions (mismatches) was 5.1%, 3.4% and 4.1%, respectively (**Fig. 1c**).

Four classes of canonical SVs (DEL, INS, DUP and INV) with a length of at least 50-bp were detected. To obtain reliable SVs, we used three SV callers, Sniffles^15^, NanoVar^16^ and NanoSV^17^, all of which were specifically designed for SV detection from LRS (**Supplementary Fig. 2a**). We retained the SVs identified by at least two callers (**Fig. 1d**), which could effectively reduce the false positive of SV detection, particularly for sequencing data with lower depth (**Supplementary Fig. 2b** and **Methods**). In addition, we applied three filtering steps that removed an average of 1,331 SVs per sample to further reduce unreliable SVs (**Fig. 1e**, **Supplementary Table 3** and **Methods**). Finally, we identified 18,489 high-confidence SVs per sample, ranging from 15,439 to 22,505 (**Fig. 1f** and **Supplementary Table 3**). The numbers of SVs followed an approximately normal distribution (**Supplementary Fig. 2c**). DELs and INSs were predominant, and each sample contained an average of 8,215 DELs (44.4%), 9,942 INSs (53.8%), 258 DUPs (1.4%) and 74 INVs (0.4%) (**Fig. 1f**). A balanced number between DEL and INS was also observed in the previously LRS-based SV studies^8,10^, and slightly higher ratio of INSs than DELs may be due to the DEL bias of GRCh38^18^.

We estimated the relationship between the SV number and sequencing depth. It was observed that the number of SVs just slightly increased when the depth was more than 15-fold (**Supplementary Fig. 2b**). The number of SVs was around 19,070 per sample at 15-fold depth and that increased to around 20,378 at 40-fold depth (**Supplementary Fig. 2b**), suggesting that 15-fold sequencing was effective to detect SVs for a population-scale study.

It is known that LRS technology such as ONT with high sequence error more likely leads to mis-mapping against the reference genome and therefore causes higher false discovery of SVs. To estimate the false discovery rate (FDR) of SVs using our SV identification strategy, SVs detected from a 15-fold ONT dataset (HG002, the child) were validated by a PacBio high-fidelity (HiFi) dataset from a parent-offspring trio in Genome in a Bottle (GIAB), whose base accuracy was up to 99.8%^19^. Out of the 18,737 detected SVs for the HG002 dataset, the overall FDR was 3.2%, illustrating the reliability of SVs detected using our SV identification strategy from ONT reads with 15-fold depth (**Methods**). INVs are generally enriched for false positives. To estimate FDR for INVs, we further manually investigated the strand-specific alignment of long-read for the INV region using Integrative Genomics Viewer (IGV)^20^. We checked 75 INVs and detected four false positives with an estimated FDR of 5.3%.

### Comparison with published SV datasets

We merged the SVs detected from all the samples for each SV type and constructed a set of 132,312 non-redundant SVs, comprising 67,405 DELs, 60,182 INSs, 3,956 DUPs and 769 INVs (**Fig. 2a**). We compared our data with four previously published datasets generated using either NGS or TGS platforms (**Fig. 2b**). To note, the LRS15 set was the only one generated using the TGS platform (PacBio) from 15 individuals^8^, which had the largest overlaps with our data (38,963) and had the smallest unique SV set of its own (60,641). For other three datasets generated using the NGS platform, there were 30,783, 24,741 and 24,472 SVs that overlapped with the Database of Genomic Variants (DGV)^21^, Genome Aggregation Database (gnomAD)^22^ and the Human Genome Diversity panel (HGD)^23^, respectively. In total, 70,471 (53.3%) SVs from our data have not been previously reported. We further examined the recovery of SVs in the previous datasets for each SV type. It was notable that although the total numbers of INSs and DELs were similar in our dataset, the number of recovered INSs with the LRS-based study, LRS15, was much larger than the SRS-based datasets, illustrating that LRS technology is particularly efficient to detect INSs (**Fig. 2c**).

**Figure 2.**
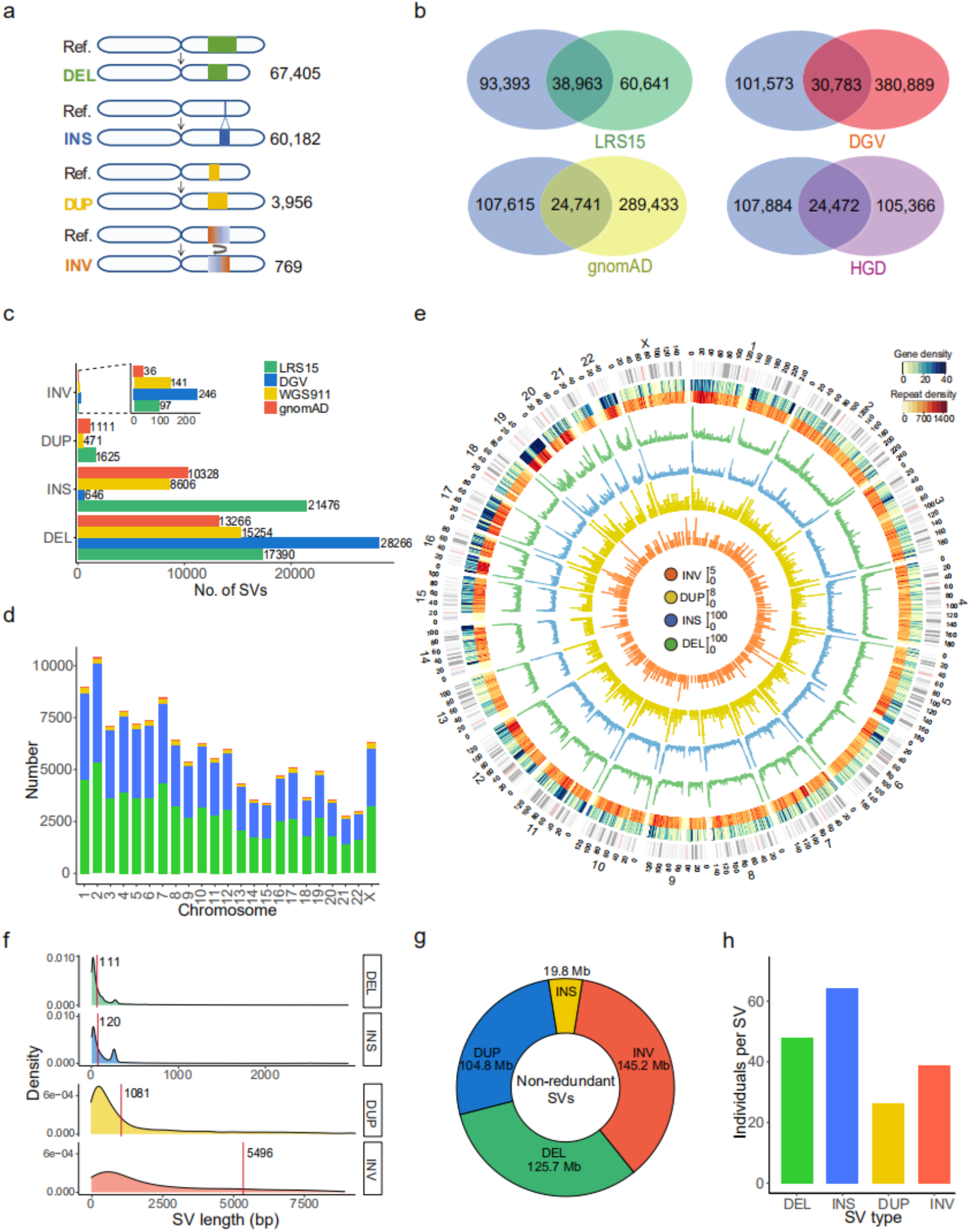
Properties of non-redundant SVs for each SV type. **a**, No. of non-redundant SVs of all individuals. **b**, Overlaps of SVs between our study (blue ellipse) and the previously published datasets. **c**, Overlaps of SV number between our study and the previously published datasets for each SV type. The number on the bar graph indicates the actual number of SVs. **d**, Total No. of non-redundant SVs for each SV type. **e**, No. of genes, repeats and SVs within 500 kb non-overlapping window across chromosomes. The two outer circles denote the distribution of genes and density of repeats, followed by distributions of DEL (green), INS (blue), DUP (yellow) and INV (orange). **f**, Length distribution for each SV type. Red line indicates the median length of each SV type. **g**, Total length of non-redundant SVs for each SV type across chromosomes. **h**, The average No. of individuals for merged non-redundant SVs of each type.

### Genomic features of SVs

SVs were nonrandomly distributed across the chromosome and the number of SVs significantly correlated with chromosome length (*R* = 0.92, *P* = 8.3×10^−10^, Pearson correlation test) (**Fig. 2d**). The number generally increased at the ends of chromosome arms, particularly for DELs, INSs and DUPs (**Fig. 2e**). The sub-telomeric bias of the long arms of chromosomes was higher compared to that of short arms (**Supplementary Fig. 3**), which was in accord with the pattern detected by an LRS-based SV study using a PacBio platform^8^.

We observed that the median lengths of INSs and DELs were 111 bp and 120 bp, respectively, which was significantly shorter than that of DUPs (1,081 bp) and INVs (5,496 bp) (**Fig. 2f**). The longer length of DUPs and INVs was confirmed by our PacBio HiFi datasets as well as the PacBio HiFi datasets of the trio from GIAB, indicating that the observation was not the ONT platform specific. The numbers of DELs and INSs rapidly decreased as their length increased. There were two clear peaks at sizes around 300 bp and 6 kilobases (kb) for both DELs and INSs (**Supplementary Fig. 4**), corresponding to short interspersed nuclear elements (SINEs) and long interspersed nuclear elements (LINEs)^8,10^.

The total length of non-redundant SVs was 395.6 Mb, representing approximately 13.2% of the human reference genome, including 125.7 Mb of DELs, 19.8 Mb of INSs, 104.8 Mb of DUPs and 145.2 Mb of INVs (**Fig. 2g**). On average, SVs affected 23.0 Mb (around 0.8%) of the genome per individual (**Supplementary Table 4**), where the average lengths of DELs and INSs were 7.2 Mb (31.2% of the total SV length) and 3.7 Mb (15.9%), respectively. Despite their lower number, INVs (9.6 Mb, 41.7%) and DUPs (2.5 Mb, 11.1%) contributed equivalently to the total SV length due to their considerably longer length. Same INSs were more common in individuals compared to those of other types, which may be partly due to the DELs bias of GRCh38 or purification selection of INSs in Chinese population (**Fig. 2h**). 75.4% of SVs contained repeat sequences (**Supplementary Table 5**), which was consistent with the knowledge that SVs tended to occur in segments with more repetitive sequences^1^. Among all of these repeats, VNTRs (25.1% of SVs) and SINEs (19.7%) were predominant (**Supplementary Table 5**).

### Allele frequencies of SVs

Our datasets offer us an opportunity to identify SVs with a low frequency in a population. We grouped the SVs into four categories based on their allele frequencies (AF): singleton (allele count = 1), rare (allele count >1 and AF ≤ 0.01), low (0.01 < AF ≤ 0.05) and common (AF > 0.05). Singletons (56,239) represented 42.5% of the total identified SVs (**Fig. 3a** and **Supplementary Fig. 5)**. Additionally, there were 28,925 rare (21.9%), 14,296 low (10.8%) and 32,852 common (24.8%) SVs. Among the common SVs, 1,264 (3.9%) were shared in all samples. The lower AF values of an identified SV, the larger proportion of novel SVs that were not previously reported (**Fig. 3a**). Specifically, 75.1% of singleton SVs were novel, which was similar to the percentage of novel singleton SNVs (72.8%) in Koreans^24^. In contrast, 14.9% of common SVs were not previously reported.

**Figure 3.**
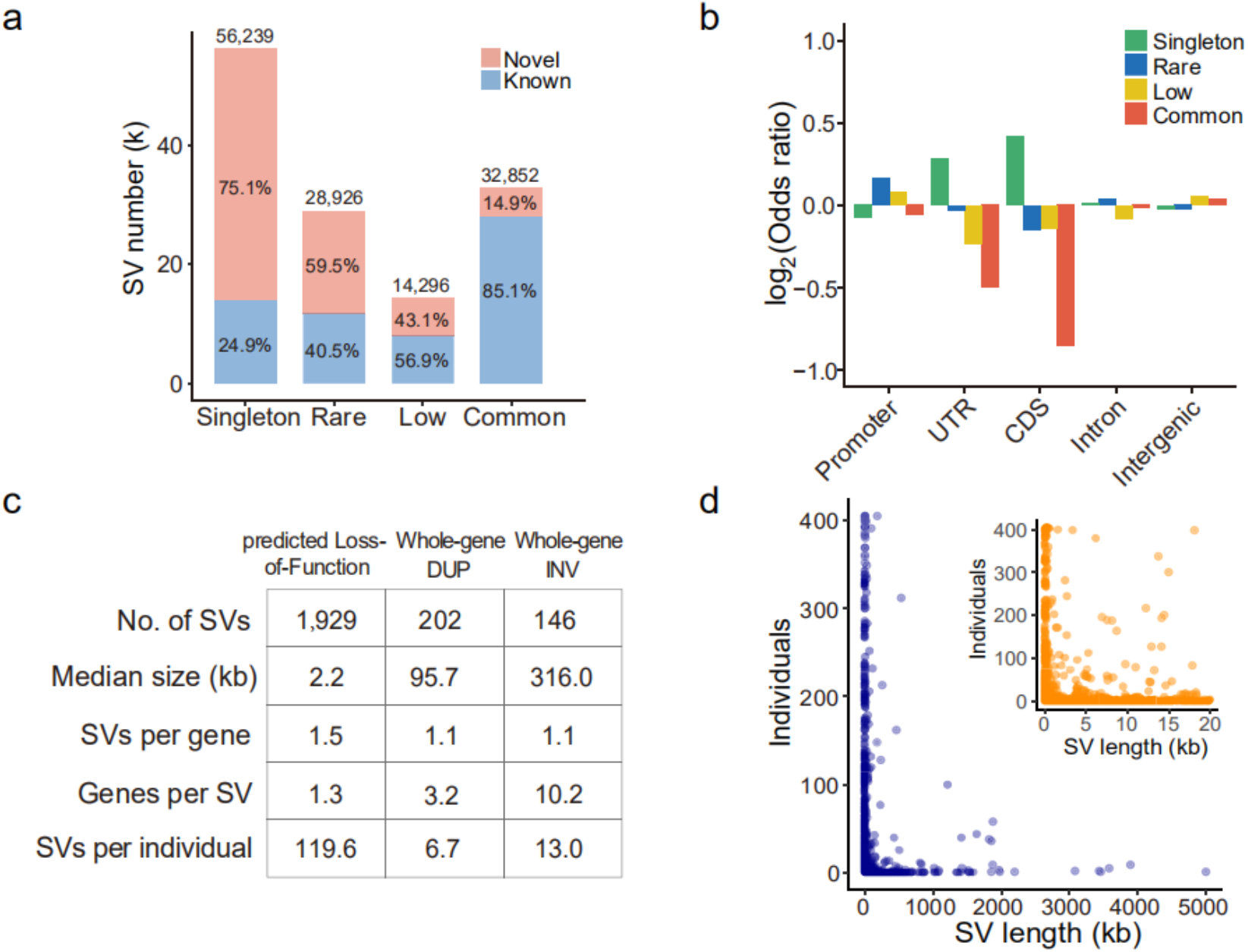
Allele frequency of SVs and functional annotation. **a**, No. of known and novel SVs for each SV category based on the variant allele frequency (AF): singleton (allele count = 1), rare (allele count >1 and AF ≤ 0.01), low (AF > 0.01 but ≤ 0.05) and common (AF > 0.05). **b**, Enrichment analysis of genomic location of SVs for each category. **c**, Statistics of predicted Loss-of-Function (pLoF) SV, whole-gene DUP (WDUP) and whole-gene INV (WINV). **d**, Individual No. versus SV length for pLoF SVs. The blue figure shows SV length from 1-5000 kb while the orange figure shows that from 0-20 kb.

Singleton SVs are prone to false positives compared with other categories because they are detected from one sample. To validate the accuracy of singletons, we designed the primers for 154 randomly selected singleton DELs and INSs from 20 samples and validated 145 by PCR experiments (FDR = 5.8%). (**Supplementary Table 6**). In addition, we sequenced four samples in our study using the Pacbio HiFi (average depth of 9.35-fold, **Methods**). There were 510 singletons detected from the ONT reads for these four samples. And we found that 32 SVs were false positive based on the validation of PacBio HiFi reads and manual investigation by IGV (FDR = 6.3%).

To estimate how much SV spectrum in Chinese population has been detected in our study, we assessed the SV numbers with different categories as the number of samples increased through multiple sampling. Relatively stable number of the common and low SVs indicated that the samples used in this study was sufficient to characterize these SVs **(Supplementary Fig. 6**). The continued increasing trends of the singleton and rare SVs suggested that a larger sample size was needed to sufficiently detect SVs with an AF ≤ 0.01.

### Functional relevance of SVs

To explore their potential functions, we annotated SVs based on their genomic location, including coding sequence (CDS), untranslated region (UTR), promoter and intron. A substantial amount (37.6%) of SVs were located in introns, while 1.0%, 0.9% and 1.7% of SVs were located in the promoter, UTR and CDS, respectively (**Table 1**). Among all the SVs located in the UTR and CDS, singletons were significantly enriched compared with the other categories (*P* = 1.1 ×10^−4^ for singletons in UTR and 5.8 ×10^−16^ for singletons in CDS, Fisher’s exact test, **Fig. 3b**), suggesting that singleton SVs were more likely to have genetic functions.

**Table 1.** Gene Annotation for SVs in Each SV Category.

| Category | Promoter (%) | UTR (%) | CDS (%) | Intron (%) | Intergenic (%) | All |
| --- | --- | --- | --- | --- | --- | --- |
| Singleton | 542 (1.0) | 594 (1.1) | 1,271 (2.3) | 21,311 (37.9) | 32,521 (57.8) | 56,239 |
| Rare | 329 (1.1) | 245 (0.8) | 441 (1.5) | 11,204 (38.7) | 16,706 (57.8) | 28,925 |
| Low | 154 (1.1) | 105 (0.7) | 219 (1.5) | 5,076 (35.5) | 8,742 (61.1) | 14,296 |
| Common | 320 (1.0) | 201 (0.6) | 303 (0.9) | 12,208 (37.2) | 19,820 (60.3) | 32,852 |
| All | 1,345 (1.0) | 1,145 (0.9) | 2,234 (1.7) | 49,799 (37.6) | 77,789 (58.8) | 132,312 |

Number and percent of SVs affecting the promoter (1 kb upstream of gene), untranslated region (UTR), coding sequence region (CDS) and intron of protein coding gene. SVs that were not intersected with gene were annotated as intergenic region.

We further classified the SVs that interacted with CDS into three subgroups according to their breakpoint locations: predicted loss-of-function (pLoF), whole-gene duplication (WDUP) and whole-gene inversion (WINV) (**Methods**). While pLoFs delete coding nucleotides or alter open-reading frames, WDUPs generally cause a copy-gain of an entire gene, and WINVs regulate gene expression through affecting the position and order of upstream enhancers and genes^25^. We annotated a total of 2,277 SVs affecting the coding regions of 3,176 distinct genes, including 1,929 pLoF SVs, affecting the CDS of 1,681 genes, as well as 202 WDUPs and 146 WINVs, covering 581 and 1,331 genes, respectively (**Fig. 3c**). Interestingly, Gene Ontology (GO) analysis revealed that there were 38 genes interested with pLoF SVs that were significantly enriched in “immunoglobulin receptor binding” (odds ratio = 5.7, adjusted P value = 7.2×10-18, Benjamini-Hochberg corrected, **Supplementary Fig. 7**).

On average, individuals carry 2.7 and 2.9 pLoF SVs for singleton and rare categories, respectively. More than half of pLoF SVs (57.6%) were singletons, and the median length of all pLoF SVs was 2.2 kb (**Fig. 3d**). The median length of common pLoF SVs was just 251 bp, while that of singleton pLoF SVs was up to 4,480 bp (**Supplementary Fig. 8**). Longer SVs in the singletons may be more likely to disrupt gene function.

### Phenotypic and clinical impacts of SVs

In order to better understand how pLoF SVs impact clinical phenotypes and diseases, we annotated these SVs and their associated genes using the Genome-Wide Association Studies (GWAS) catalog^26^, Online Mendelian Inheritance in Man (OMIM)^27^ and Catalogue Of Somatic Mutations In Cancer (COSMIC)^28^. For a total number of 1,929 pLoF SVs, 1,231 (63.8%) were intersected with genes cataloged in above databases (**Supplementary Table 7**). Among the 1,231 SVs, 58.1% to 60.2% belonged to the singleton category (**Fig. 4a**), which was consistent with our previous enrichment analysis showing that singletons were more likely to be functional (**Fig. 3b**). At the gene level, all of the 1,929 pLoF SVs intersected with 1,681 distinct genes, where 957 genes (56.9%) were annotated in the three databases (**Fig. 4b** and **Supplementary Table 7**).

**Figure 4.**
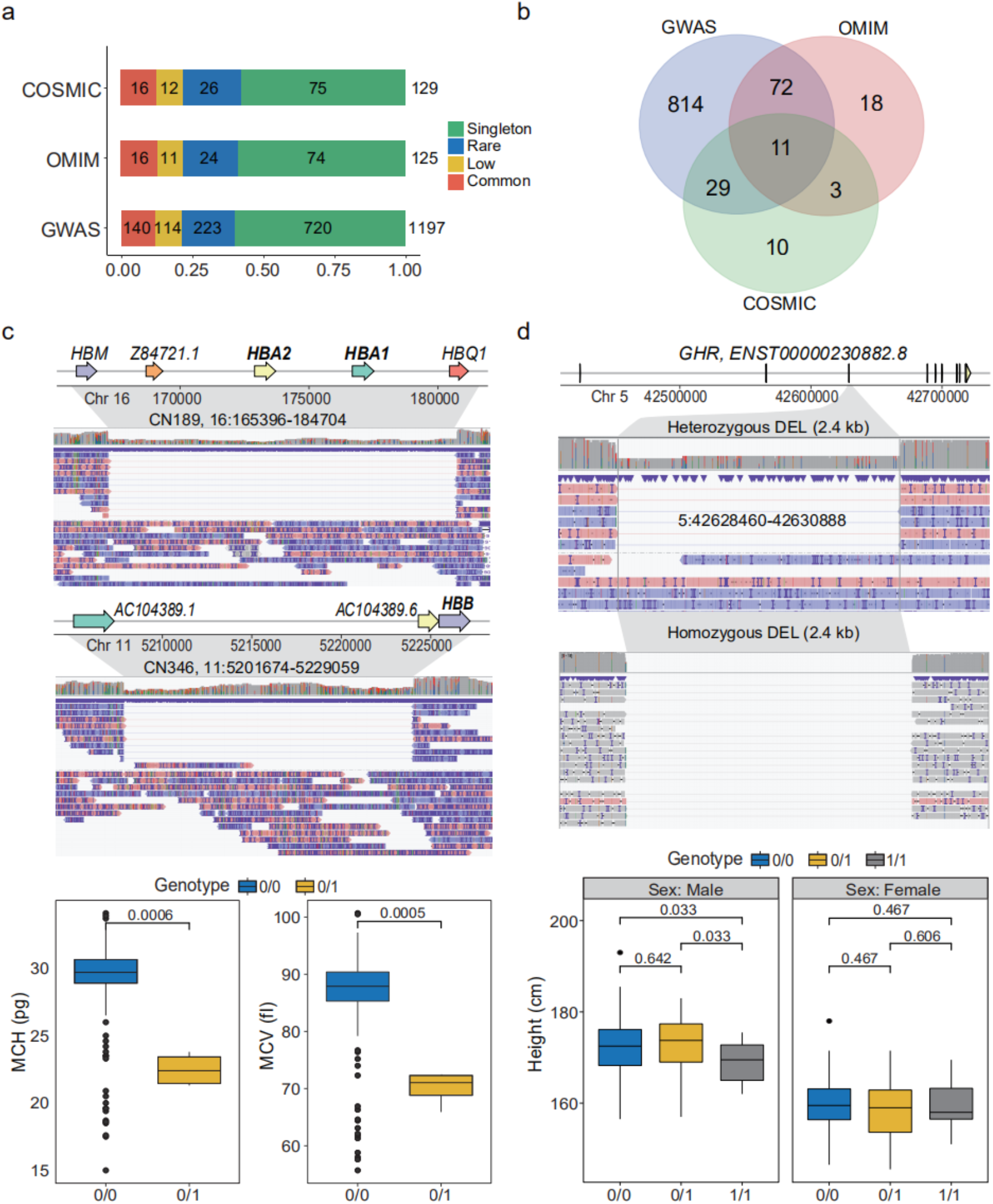
pLoF SVs associated with phenotypes and diseases. **a**, No. of pLoF SVs with reported genes in GWAS, OMIM and COSMIC for each SV category. **b**, No. of genes associated with pLoF SVs. **c**, Example of SVs affecting *HBB* and *HBA1*/*HBA2* which are associated with anemia. Up: IGV screenshot of a 19.3 kb heterozygous DEL covering both *HBA1* and *HBA2*; Middle: IGV screenshot of a 27.4 kb heterozygous DEL covering *HBB*; Bottom: mean corpuscular hemoglobin (MCH) and mean corpuscular volume (MCV) values of four individuals containing a 19.3 kb DLE (3 individuals) and a 27.4 kb DEL (1 individual) and the others. **d**, Example of SVs affecting *GHR* which is associated with height of male adults. Up: IGV screenshot of a 2.4 kb heterozygous DEL completely covering the third exon of *GHR*; Middle: IGV screenshot of the 2.4 kb homozygous DEL. The *p* values were derived from multiple testing correction. For **c** and **d**, “0/0”: homozygous allele same as the reference; “0/1”: heterozygous variant; “1/1”: homozygous variant.

Many phenotypically and clinically relevant SVs could be discovered from our dataset. For example, we detected plausible causal variants associated with anemia which has not been reported. We found a heterozygous rare DEL of 19.3 kb in three individuals, covering the genes Hemoglobin Subunit Alpha 1 and 2 (*HBA1* and *HBA2*), whose dysfunctions are known to cause α-thalassemia^29^ (**Fig. 4c**). In addition, one individual had a heterozygous DEL of 27.4 kb, containing gene Hemoglobin Subunit Beta (*HBB*), whose dysfunction is known to cause serious hemoglobinopathies, such as sickle cell anemia and β-thalassemia (**Fig. 4c**)^30^. As expected, the mean corpuscular hemoglobin values (MCH, ranging from 21.3 to 23.8 pg) and the mean corpuscular volume values (MCV, ranging from 65.9 to 72.3 fL) of these four individuals carrying the aforementioned heterozygous DELs were significantly lower than individuals with the reference allele in our study (*P* = 0.0006 and 0.0005 for MCH and MCV, respectively, *t*-test corrected by FDR, **Fig. 4c**).

Besides rare SVs, we could also detect common SVs that were associated with various phenotypes. We observed a DEL of 2.4 kb, which existed in 35 homozygous (15 males and 20 females) and 67 heterozygous (35 males and 32 females) carriers in our study, covering the complete region of the third exon of *GHR* (Growth Hormone Receptor), whose missense mutation is known to cause short stature and dwarfism (**Fig. 4d**)^31^. Interestingly, the homozygous carriers were significantly shorter than heterozygous carriers and people with homozygous reference allele for males (168.9 vs. 172.9 (cm) and 168.9 vs. 172.4 (cm), *P* = 0.033 for both, *t-*test corrected by FDR, **Fig. 4d**), suggesting that the DEL acted as recessive allele for this gene. In contrast, no significant difference was observed between these three genotypes in females. The growth hormone (GH)–insulin-like growth factor (IGF)-I axis is the principle endocrine system regulating linear growth in children^32^. This sex different phenotype could be caused by the cross-talk between the GH/IGF axis and sex hormones, where pituitary GH secretion regulates many sex-dependent genes in the liver^33^, and the pituitary is the major source of IGF that determines body height, in a sex-dependent manner^34^.

We further conducted Genome-wide association study (GWAS) for the clinical phenotypes based on genotyped SVs with minor allele frequency (MAF) > 0.05. We found that 25 SVs were significantly associated with 14 phenotypes on 13 chromosomes (*P* < 1.7×10^−6^, the Bonferroni-corrected significance threshold, **Supplementary Table 8**). Of which, 9 SVs were in the introns, and the remaining 16 SVs were in the intergenic regions. One example is that a 114 bp DEL with MAF of 0.10 on Chromosome 5 was significantly associated with urinary crystal (XTAL). This DEL located in the intron of *SLC9A3* (sodium/hydrogen exchanger, isoform 3), that was previously found to be associated with ammonia metabolism which regulated acidic or alkaline in urine^35^ and thus affected the calcium oxalate (CaOx) crystallization^36^.

### Population evolution of SVs

Previous population genetics studies have shown the genetic differences between northern and southern Chinese using SNP arrays and NGS-based WGS^11,37^. We herein explored the population genetic properties of SVs between these two regional groups. Principal component analysis (PCA) showed the clear genetic diversity between the two groups, revealing that population structures were consistent with self-reported ancestry (**Fig. 5a**). We further estimated the differentiation between them. The average value of the fixation index (*F*_st_) was 0.0032, which was slightly higher than the value estimated by SNVs (*F*_st_ = 0.0015) in a previous study^11^, implying a higher divergence between the two sub-populations in this study. We observed 15 significant signals (*F*_st_ > 0.066, **Supplementary Table 9**) across the genome, which were clustered into eight peaks on Chromosomes 1, 2, 3, 6, 10, 12, 14 and 19 (**Fig. 5b**). Among the 15 signals, four SVs were located within four genes, respectively, namely *HCG4B*, *IGHG3*, *MUC4* and *SLC1A7* (**Fig. 5c** and **Supplementary Table 9**). The top two *F*_st_ clusters were located in the major histocompatibility complex (MHC) region at Chromosome 6 and in the immunoglobulin heavy (IGH) cluster locus at Chromosome 14. In the MHC region, significantly differentiated SVs located in the exons of *HCG4B* (HLA complex group 4B) and intergenic regions of *HLA-K* and *HLA-U* while MHC was a known site of extreme genetic diversity across humanity and was reported to be under selection in East Asian population^38^. (**Fig. 5c** and **Supplementary Table 9**). It is also notable that although there were seven SVs were significantly differentiated on the IGH cluster loci, no haplotype block was observed (**Fig. 5d**), which illustrated the genetic diversity among individuals, suggesting that accumulation and combination of different genotypes of IGH genes might be associated with the immunity adaption to diverse environments. In addition, a 477 bp DEL was observed in intron of *SLC1A7*, which is a solute carrier family member and has L-glutamate transmembrane transporter activity and was previously reported to be the immune-associated prognosis signature for hepatocellular carcinoma^39^. The differentiation of SVs in the immunity associated regions might be due to genetic drift and long-term expose to diverse environments for sub-populations of Chinese.

**Figure 5.**
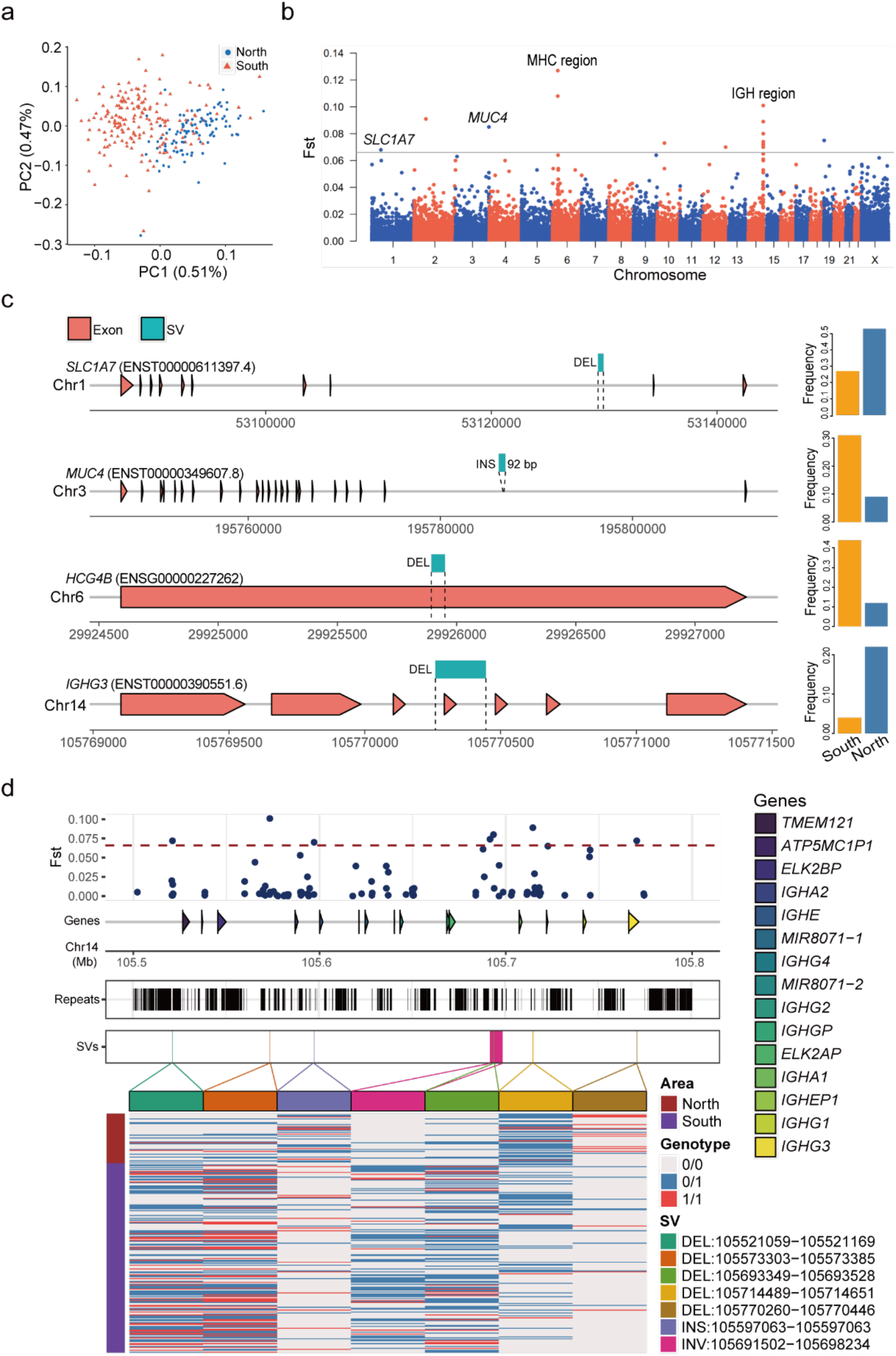
Genetic differentiation of SVs between sub-populations of Chinese. **a**, PCA of the two sub-populations: northern and southern Chinese. The values in parenthesis indicate the genetic variations explained by first two PCs. **b**, *F*_st_ value between the two sub-populations. The gray horizontal line indicates the *F*_st_ cutoff of > 0.066 based on permutation. **c**, Four significantly differentiated SVs located in genes. Barplots on the right panel showing the frequency of each SV between northern and southern sub-populations. **d**, Differential SVs clumped to IGH region in chromosome 14 and the genotype patterns between the two sub-populations.

## Discussion

This study presented one of the largest LRS-based genomic datasets to date. We generated reliable reference sets for SVs and identified a mean of 18,489 high-confidence SVs per Chinese genome that affected 23.0 Mb of the genome. Our SV datasets was much larger than the ones from SRS-based studies, where an average of 4,442 and 8,202 SVs per human genome were detected^25,40^. Out of 132,312 non-redundant SVs described here, 53.3% were previously unreported. The high number of novel SVs in our study might be due to: (1) the methodological improvements of detecting SVs using LRS technology; (2) the large number of samples in our study; and (3) the inclusion of Chinese population, which has been lowly represented in previous studies.

LRS technology such as ONT with relatively high sequence error rates more likely leads to mis-mapping against the reference genome and may cause unreliable SV detection. We used multi-algorithm ensemble approach and stringent filtering strategy to improve SV detection. In addition, we performed orthogonal validation using PacBio high-fidelity (HiFi) sequencing and PCR experiments. The overall FDR of all the SVs was around 3.2% and that of singleton SVs was around 6%. Compared to a prior study where an average number of 22,755 SVs per genome was detected by an SV caller, SMRT-SV^8^, our study detected a lower number of SVs per genome (18,489), mainly because we used multiple SV callers and stringent filtering processes. When we applied one caller, such as Sniffles or NanoVar, to detect the SVs, we could detect an average of 20,186 or 31,252 SVs per sample, respectively. When estimating the relationship between the detected SV number and sequencing depth, we found that the number of SVs just became stable when the depth was more than 15-fold. Our analysis suggested that our strategy was effective to detect SVs for a population-scale study.

Our datasets enable us to explore SVs with a low frequency in the population. We provided several lines of evidence that singleton and rare SVs were more likely to be functional. In particularly, pLoF SVs that altered coding regions and affected clinical phenotypes could be rare or even singleton in the population, such as long DELs covering the whole genes of *HBA1*/*HBA2* and *HBB*. Indeed, a recent study also discovered rare LoF variants in 26 genes, derived from whole–exome sequencing, were significantly associated with phenotypes^41^.

With the rich phenotypic and clinical measurements, our study presents a key step for establishing a regional reference genome and provides a broad prospect to improve the interpretation of clinical genetics in Chinese population. To date, many diseases, such as rare diseases and neurodevelopment-related diseases, have been validated to be caused by SVs^42,43^. However, it is still a difficult task to find pathogenic SVs, even though conducting a whole-genome scan using LRS approaches, as the effects of SVs still remain largely unknown, particularly for those in noncoding regions. Our SV dataset constructed from a large Chinese population could help future LRS-based genomic studies to narrow down candidates for pathogenic variants.

In summary, given that the current human reference genome and population genomics have a substantially large number of uncovered SVs, this study presents an important effort to fill this knowledge gap and provides us the opportunity to detect novel SVs associated with phenotypes, diseases and evolution.

### Ethics approval and consent to participate

This research was approved by the Ethics Committee of Zhongshan Ophthalmic Center, Sun Yat-sen University (2019KYPJ111) and West China Hospital, Sichuan University (2018-120). Clinical data and biological specimens were obtained from the individuals with written informed consent.

## Data availability

SVs of 405 individuals have been stored in VCF format and deposited into the Genome Variation Map (GVM) database in Beijing Institute of Genomics (BIG) Data Center (https://bigd.big.ac.cn/gsa/index.jsp). The codes of data analysis and PCR validations are available at https://github.com/xie-lab/PGC. All the data and codes will be publicly available.

## Acknowledgements

Dr. Rong Ju provided helpful suggestion. We thank the support of the Center for Precision Medicine at Sun Yat-sen University. We also thank BioMarker Inc. for providing the ONT sequencing platform and Annoroad Gene Technology company for providing the PacBio sequencing platform.

## Author contributions

Z.X conceived and supervised the study. ZK.W, Z.X, Tao L and YZ. L designed the study. ZK.W, Tong L, ZH.J and Z.X analyzed the data. LS.Z, CB.X, S.OY and Tao L organized and executed the clinical study. JQ.Y conducted PCR experiments. ZK.W, Tong L and ZH.J prepared the manuscript. ZK.W, Z.X and Tong L wrote the manuscript. All authors read and approved the manuscript.

## Methods

### Sample information

A total of 405 individuals were enrolled in this study (206 males and 199 females) with age varying from 22 to 81 years old (**Fig. 1a**). These individuals came from 18 provinces in China according to their self-reported original province. Northern and southern Chinese were distinguished based on Qinling Mountain-Huaihe River Line. Among them, 329 individuals were recruited at the Health Service Center of Sun Yat-sen University Cancer Center, where 68 clinical phenotypes from 327 individuals were collected. Additional 76 individuals from the West China Hospital of Sichuan University were included in this study. Written informed consent was obtained for all the individuals.

### Phenotype collection

Anthropometric measurements, including height, weight and body mass index (BMI), were obtained from an automatic electronic meter (SECA GM-1000, Seoul, Korea). Blood tests were processed by a hematology automated analyzer (SYSMEX XE-2100, Kobe, Japan). Urine tests were determined using an automated urine chemistry analyzer (ARKRAY4030, Tokyo, Japan) and a urinary tract infection analyzer (SYSMEX UF-1000i, Kobe, Japan). Biochemical detection was performed by an automatic modular analyzer (Cobas C701, Basel, Switzerland). Tumor markers such as alpha fetoprotein (AFP) and carcinoembryonic antigen (CEA) were measured by an immunology modular analyzer (Cobas 8000 e602, Basel, Switzerland).

### Library construction and long-read sequencing

High molecular weight genomic DNA of each individual was extracted from peripheral blood leukocytes using HiPure Tissue & Blood DNA Kit (D3018-03, Angen). For the Nanopore sequencing, DNA repair, end repair and adapter ligation were conducted during library preparation, and 2 μg DNA was Fragmented by g-TUBE (Covaris). DNA repair was performed using NEBNext FFPE DNA Repair Mix (M6630L, NEB). End repair was performed using NEBNext Ultra II End Repair/dA-Tailing Module (E7546L, NEB). Adapter ligation was performed using NEBNext Blunt/TA Ligase Master Mix (M0367L NEB) and Ligation Sequencing Kit 1D (SQK-LSK109, Oxford Nanopore Technologies). DNA was purified between each step using Agencourt AMPure XP beads (A63882, Beckman Coulter). DNA was quantified via a Qubit Fluorometer 2.0 (ThermoFisher Scientific, Waltham, MA). We carried out long-read sequencing using a PromethION sequencer and 1D flow cell with protein pore R9.4.1 1D chemistry according to the manufacturer’s instructions. Reads were base-called in batches by guppy v3.2.8 using the default parameters during sequencing.

For the PacBio sequencing, the integrity of the DNA was determined with the Agilent 4200 Bioanalyzer (Agilent Technologies, Palo Alto, California). Eight micrograms of genomic DNA were sheared using g-Tubes (Covaris), and concentrated with AMPure PB magnetic beads. Each SMRT bell library was constructed using the Pacific Biosciences SMRTbell template prep kit 1.0. The constructed library was size-selected by Sage ELF for molecules 11~15 kb, followed by primer annealing and the binding of SMRT bell templates to polymerases with the DNA Polymerase Binding Kit (Pacific Bioscience). Sequencing was carried out on the Pacific Bioscience Sequel II platform for 30 hours.

### Read quality control and mapping

We detected lower quality bases at two ends of reads by NanoQC^1^ v0.8.1 and trimmed 30 bases of start and 20 bases of end for each raw read using NanoFilt^1^ v2.2.0 due to their lower quality. We kept the reads with length longer than 500 bp and mean quality higher than seven for downstream analysis. The statistics of length and quality value of clean reads was performed using NanoPlot^1^ v1.20.0. The clean reads were then aligned to the primary assembly of human reference genome GRCh38 using minimap2^2^ v2.15-r905 with the recommended option for ONT reads (−x map-ont) and the additional parameters “--MD -a”. Aligned files with SAM format were converted to BAM format and then sorted using SAMtools^3^ v1.9. Summary of aligned information for BAM file was conducted by command “stat” of SAMtools and region depth of aligned file was estimated by mosdepth^4^ v0.2.5. The read base mapping rate and sequencing error rate were estimated with the method previously described^5^. Specifically, we aligned the cleaned reads to GRCh38 and estimated the rates of substitutions, insertions and deletions based on the mapping result, excluding secondary alignments and the soft-clipped sequences.

### Detection of high-confidence SVs

To obtain high-confidence SVs, we employed multiple tools to call SVs and conducted a series of filtering steps (**Supplementary Fig. 2a**). Sniffles^6^ v1.0.10, NanoVar^7^ v1.3.6 and NanoSV^8^ v1.2.4, which were SV callers specifically designed for long-read, were used to detect SVs. Sniffles was used with parameters “--min_support 2 --min_length 50 --num_reads_report −1 -- min_seq_size 500 --genotype --report_BND --report_seq”. We set the following parameters for NanoSV: “--data_type ont --mincov 2 --minlen 50”. NanoVar had been run with the default parameters.

We merged the SV call sets of each individual derived from the above three SV callers for each type SV. We applied the Cluster Affinity Search Technique algorithm (CAST)^5,9^ to merge SVs independently for each SV type based on the variant position and length^5^. In order to facilitate the implementation of this algorithm for INSs, the end coordinate of INS was set as the sum of start coordinate and the SV length. First, we segregated all discovered SVs into non-overlapping groups. For each group, we represented SVs as nodes in a graph and drew an edge between two SVs if they had a minimum mutual overlap of at least 50% of the length. The SV merging can be modeled by a corrupted clique graph. Consequently, the merged SVs detected by at least two callers were extracted. As suggested by benchmark analysis of LRS callers in our study (Nicolas Dierckxsens et al., unpublished), Sniffles showed the most balanced performance following by NanoVar and NanoSV for ONT reads. Therefore, we prioritized the results of Sniffles, followed by NanoVar. To obtain high-quality SVs, we then conducted three steps to filter out lower quality SVs (**Supplementary Fig. 2a**). First, we extracted SVs that were supported with at least three reads. In addition, INSs and DELs with length larger than 2 Mb were discarded. While INVs and DUPs larger than 5 Mb were discarded. Furthermore, sites in the centromere region with length of 61.9 Mb in 22 autosomes and the X chromosome were removed from further analysis. We identified and discarded SVs intersected with gap regions (marked as “N”) and high depth regions (≥ 500×, estimated by mosdepth^4^ v0.2.5) with BEDTools^10^ v2.27.1. The genomic position of gap and centromere regions were downloaded via the UCSC Tablebrower^11^.

To discern the relationship between the discovery rate of SVs and sequencing depth using different SV callers, we applied the above strategy to the Nanopore (ONT) reads of HG002 from Genome in a Bottle (GIAB). First, we got the sparse sequences with different depths ranging from 8× to 40× by randomly selecting reads from the deep sequencing clean data of HG002. Then the mapping tool and SV callers coupled with the same parameters in this study were used to detected SVs for each dataset. The method “Combine” means the SV shared by at least two of three callers (**Supplementary Fig. 2b**). The ratio threshold for SV supported reads was 0.2, which was equal to 3 supported reads for 15-fold data. We found that a single caller was not sufficient to obtain high-confidence SVs, especially for depth lower than 10×. The results combined from callers with higher sensitivity (NanoSV) and precision (NanoVar), and balanced performance (Sniffles) can be used for detecting high-confidence SVs. At the same time, our stringent strategy might discard some true SVs, which contributed to the smaller number of detected SVs compared with previously study^12^. In addition, the SV number became stable when the sequencing depth was more than 15×. For Combine method, 19,070 SVs were detected at depth of 15×, which was 93.6% of the total number (20,378) at depth of 40× (**Supplementary Fig. 2b**), indicating that the median depth (15×) sequencing data was a cost-effective method for genetic research at a population-scale.

### Non-redundant SVs and genotypes in the population

A large proportion of SVs were carried by multiple individuals because of their genetic similarity within population. To remove redundancies, we merged SVs of all the individuals using CAST algorithm. Any region that frequently occurred across the population was selected to represent the non-redundant SVs. SVs were genotyped based on the variant allele balance (VAB). The genotype of individual was assigned as “0/0” if VAB ≤ 0.2, and genotype was “0/1” and “1/1” for 0.2 < VAB ≤ 0.8 and VAB > 0.8, respectively^13^. The threshold was same to prior study by Pedersen et al.^13^and similar to that of vg toolkit (0.14 for default) when applying graph genotyping^14^.

After genotyping, the merged SVs were classified with four categories based on the variant allele frequency (AF): singleton (allele count = 1), rare (allele count >1 and AF ≤ 0.01), low (0.01 < AF ≤ 0.05) and common (AF > 0.05). To estimate the relationship between non-redundant SV number for different categories and sample size, we merged SVs randomly sampled from 100 to 405 samples while setting step size as 3, and repeated four times for each step. Then we calculated SV number for each category and regarded the average of four times as the estimate value of each step. We observed that the number of common SVs in population was relatively stable (**Supplementary Fig. 6**). As samples increased, the number of low SVs decreased and rare SVs increased, and the steps appeared when the sample number was a multiply of 50 due to the same integer threshold in this period. However, the total number of low and rare SVs increased with similar trend of singletons.

### False discovery rate (FDR) of detected SVs

In order to estimate the false discovery rate (FDR) for the detected SVs, we applied the strategy used in this study to the published dataset comprising of a parent-offspring trio with ONT and PacBio high-fidelity (HiFi) reads (accuracy > 99%). The depths of PacBio HiFi reads for HG002, HG003 and HG004 were 19.5×, 21.9×, and 21.6×, respectively. Simultaneously, we randomly selected 15.1× depth sequences from cleaned ONT reads for HG002 (child). After detecting SVs for each dataset via the same strategy as stated before, we compared SVs from HG002 with ONT reads to those of the trio with HiFi reads. For the 18,737 SVs detected by ONT reads, there were 1,165 (6.2%) SVs not detected by HiFi reads of the trio. After manual investigation of IGV snapshot of HG002 with ONT reads and the corresponding PacBio HiFi reads, we finally found 608 false positive SVs in HG002 with ONT reads, and the FDR of SV detection was 3.2%. Among them, false positives for DEL, INS, DUP and INV were 459 (5.4%), 133 (1.3%), 11 (6.3%) and 4 (5.3%), respectively.

Singletons are known to have a higher error rate compared with the other categories because they existed in only one sample. To further orthogonally validate the accuracy of singletons uncovered in this study, we sequenced PacBio high-fidelity (HiFi) reads for CN365 (10.5×), CN366 (7.7×), CN371 (7.8×) and CN372 (11.4×) and then detected SVs applying above method. Among 510 singletons discovered by ONT reads for these samples, 32 SVs were false positive based on the validation of PacBio HiFi reads and manual curation, with an FDR of 6.3%.

Besides the orthogonal validation for singletons using PacBio HiFi data, we further validated singletons using PCR experiments. We randomly selected singletons from 20 samples with SV lengths ranging from 60 to 810 bp (average of 293 bp) (**Supplementary Table 6**) and designed the primers using BatchPrimer3^15^ to amply the SV fragments. We conducted each PCR for positive sample, followed by negative sample and purified water without DNA, which were consider as negative control. Totally, we designed primers for randomly selected 154 DELs and INSs. Among them, amplified lengths for 145 (94.2%) primers were consistent with the targets, and nine primers failed to amplify target fragments (https://github.com/xie-lab/PGC/tree/master/data/PCR). The estimated FDR of singletons was 5.8% (**Supplementary Table 6**).

### SV density of Meta-chromosome

To compute the density of SVs of chromosomes, we normalized the lengths of all 22 autosomes and X chromosome. First, we split the chromosomes into p-arm and q-arm. The value of 0 to 1 corresponded to the telomere to centromere of p-arm, and the value of 1 to 2 corresponded to the centromere to telomere of q-arm. For each arm, we set a window of 100 kb and then calculated the SV number in each overlapping window. We normalized value of each window based on the positions relative to total length of each arm.

### Comparison of non-redundant SVs to the published datasets

To assess the known and novel SVs for our non-redundant SV call set, we compared it to some published datasets, including LRS study of 15 human genomes (LRS15)^12^, Database of Genomic Variants (DGV, release 2-25-2020)^16^, Genome Aggregation Database (gnomAD v2.1)^17^ and Human Genome Diversity panel (HGD)^18^. We extracted the position relative to GRCh38 and length information for each SV. The hg38 coordinates of gnomAD was converted by LifeOver (https://genome-store.ucsc.edu/)^19^ based on the original hg37 version. Copy number variation (CNV) with copy gain and copy loss were regarded as DUP and DEL, respectively. Additionally, the mobile element insertion (MEI) in those datasets were considered as INS. The end position of INS was defined as the sum of original end and the length of INS when comparing INSs between different datasets. We excluded INSs whose insertion length was not available because both SV length and position information should be taken into account. To compare with our dataset, we conducted algorithm CAST to independently remove redundant SVs for each downloaded dataset as described in SVs merging. Intersected regions for each SV type between our study and the published datasets were conducted using BEDTools, and SVs were considered as overlapped if the reciprocal overlap was larger than 50%.

### Repeat analysis of SV sequence

In order to better evaluate the pattern of repeat sequences for SVs, we selected the sequence of the individual with longest SV length in each merged SV. Consequently, we successfully obtained 55,476, 42,912, 3,956 and 770 sequences for DEL, INS, DUP and INV, respectively (**Supplementary Table 3**). In aggregate, 103,114 (77.9% of total SVs) sequences were used for downstream analysis of repeat pattern. The repeat sequences were searched by RepeatMasker v4.0.9 (http://www.repeatmasker.org/) based on databases of Dfam^20^ v3.0 and RepBase^21^ (release 10-26-2018) with command “RepeatMasker-species human -pa threads - gff -dir output sv_seq.fa” and Tandem Repeat Finder (TRF)^22^ v4.09 with command “trf 2 7 7 80 10 50 500 -f -d -m”. Each SV was classified into the repeat family if it was occupied by more than half of the SV length. For tandem repeats, the repeat unit length ≥ 7 bp were annotated as variable number of tandem repeats (VNTR). The VNTR regions for genome reference GRCh38 were downloaded via UCSC Table brower.

### Gene features of SVs

We annotated detected SVs based on the known protein-coding gene annotation file (gtf) corresponding GRCh38 from Ensembl. We detected the intersection of SVs using BEDTools. Promoter was defined as the 1 kb region directly preceding the transcription start site of gene. We predicted Loss-of-Function (pLoF) SVs as follows: (1) DEL: overlap with at least one CDS; (2) INS: insertion directly into any CDS; (3) DUP and INV: partially overlap with at least one CDS. In addition, INV and DUP were generally long, we hence defined DUP and INV that covered the whole-gene as WDUP and WINV, respectively^23^. We did not consider WDUP and WINV as gene-disruptive SVs, although we cannot rule out the possibility that they might enhance or regulate gene expression via duplication or cis-action. In addition, we labeled SVs as UTR-disruptive if at least one breakpoint was in 5’ or 3’ UTR and this SV was not intersected with CDS. Then we labeled SVs as promoter-disruptive if at least one breakpoint was in promoter of a gene and this SV was not intersected with CDS and UTR. We labeled SVs as intron-disruptive if both breakpoints were in same gene and this SV did not meet any of the above criteria. Ultimately, the remaining SVs that were not intersected with any protein-coding gene region (including promoter) was labeled as intergenic.

### Enrichment analysis of pLoF SVs and associated genes

For enrichment analysis of each gene feature annotation of SVs, the expected value was defined as the SV number in gene feature divided by the total number of SVs in population, and the SV number in certain category of this feature divided by the total number of SVs in this category was considered as observed value. The Fisher’s exact test was conducted in R^24^ v3.5.3 (http://www.R-project.org/). To assess functions and associated pathways of pLoF SVs, we performed enrichment analysis using GSEApy v0.9.16 (https://github.com/zqfang/GSEApy). The annotation files, including GO_Molecular_Function_2018, KEGG_2019_Human, GWAS_Catalog_2019, OMIM_ Expanded, were downloaded from the Enrichr^25^ website (https://amp.pharm.mssm.edu/Enrichr). The p value was calculated with Fisher’s exact test, and multiple testing of p values were corrected by Benjamini-Hochberg method^26^.

### Population stratification and differentiation analysis

To assess the population stratification between northern or southern Chinese sub-populations, we performed principal component analysis (PCA) using EIGENSOFT^27^ v7.2.1. Previous study indicated that distinctly defined population structure was uncovered by DELs in comparison with other type SVs^18^, such as INSs or DUPs. Therefore, 56,544 DELs was used to estimate population stratification after filtering out DELs uniquely existing in southwest Chinese in this study. *F*_st_ between northern and southern Chinese sub-populations was estimated based on SVs of the individuals with self-reported ancestry information. The Hudson’s estimator *F*_st_ ((H_t_ - H_s_) / H_t_)^28^ between northern and southern Chinese was calculated. H_t_ indicates heterozygosity between subgroups, H_s_ indicates average heterozygosity within subgroups. Here H_t_ = (p_1_+p_2_)*(2-p_1_-p_2_)/2, H_s_ = p_1_*(1-p_1_) + p_2_*(1-p_2_), p_1_ and p_2_ indicate the allele frequency in two sub-groups, respectively. We used a permutation approach to estimate the significant threshold of observed *F*_st_ values. For northern Chinese, 52 individuals from provinces with high-latitude (Jilin, Liaoning, Heilongjiang, Neimenggu, Ningxia and Qinghai) were selected. To determine the significant threshold of *F*_st_, individuals (198 southern Chinese and 52 northern Chinese) from the two groups were randomly split into two sets of the original size for 1,000 times. The max *F*_st_ values across all permutations were recorded and finally arranged in descending order. The five percentile *F*_st_ value (0.066) of ranked values was the empirical genome-wide significance threshold for the overall significance level of α= 0.05^29^. Manhattan plot was performed using modified script based on qqman (https://github.com/stephenturner/qqman).

### Genotype-phenotype association analysis

The 29,510 SVs with minor allele frequency (MAF) larger than 0.05 in 327 individuals with clinical phenotypes were used for the analysis. The genome-wide association study (GWAS) was performed using PLINK^30^ v1.90b4 with linear regression under an additive genetic model for the quantitative traits, and age, sex, body mass index (BMI), and the first two principal components were included as covariates. When applying BMI GWAS, BMI itself was excluded from covariates. The association test for case-control was conducted using logistic regression module. We set the genome-wide significance threshold as 5×10^−8^, and the significance threshold was set to be 1.7×10^−6^ through Bonferroni correction (0.05/29,510)^31^.

### Visualization of SVs with long-reads

Visualization of detected SVs was performed using Integrative Genomics Viewer (IGV)^32^ v2.8.6 which was specially updated for viewing variants of long-read. For target SVs, parameter “Link supplementary alignments” was selected to clearly identify heterozygous SVs based on the split reads. For INVs, the linked long reads with different colors (red and blue) indicated different strands when aligning to reference genome.

### Statistical analysis

The statistical tests used were described throughout the article and in the figures. The one-tailed Student’s *t* test was performed to compare the clinical phenotype level between different genotypes of genes. We performed FDR correction (https://www.sdmproject.com/utilities/?show=FDR) for multiple comparisons. The enrichment analysis of singletons for different gene location was conducted by Fisher’s exact test. Benjamini-Hochberg corrected of *P* value was used for multiple test analysis. Pearson correlation coefficient was estimated for correlation analysis. All statistical tests were performed in R^24^ v3.5.3 (http://www.R-project.org/). In the boxplots, the upper and lower hinges represented the first and third quartile. The whiskers extended to the most extreme value within 1.5 times the interquartile range on either end of the distribution. The center line represented the median.

## Supplementary Figures and Tables

**Supplementary Figure 1.**
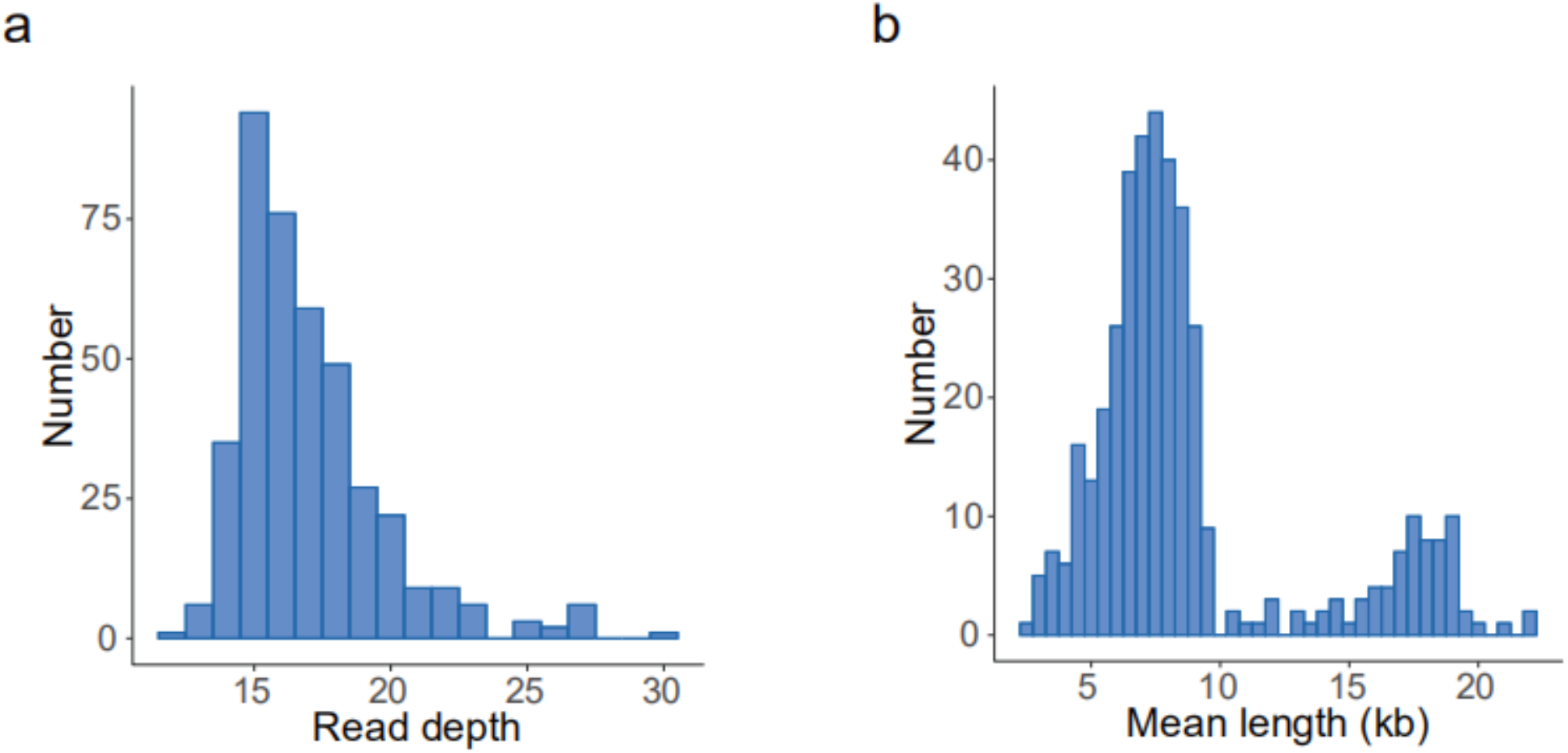
Quality control of long-reads. **a**, Read depth distribution of clean data for 405 individuals. **b**, Distribution of the average read length for 405 individuals.

**Supplementary Figure 2.**
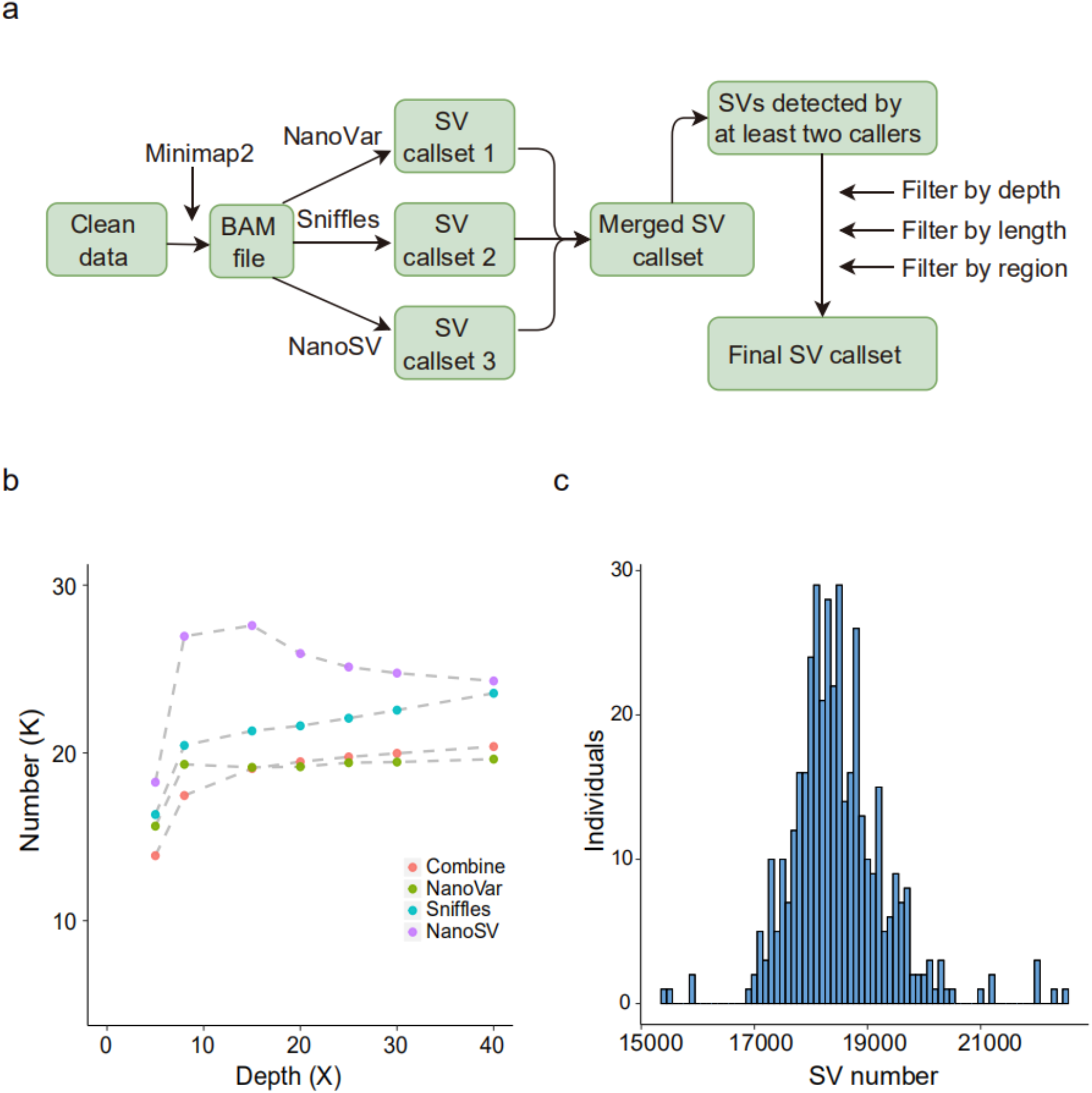
Workflow of SV calling and filtering. **a**, Workflow of SV calling and filtering for each sample. **b**, SV numbers of different callers for reads of different coverages, “Combine” means SVs detected by at least two callers. The threshold of read support of SVs is 0.2 of sequencing depth. **c**, Distribution of final high-confidence SVs of all the individuals.

**Supplementary Figure 3.**
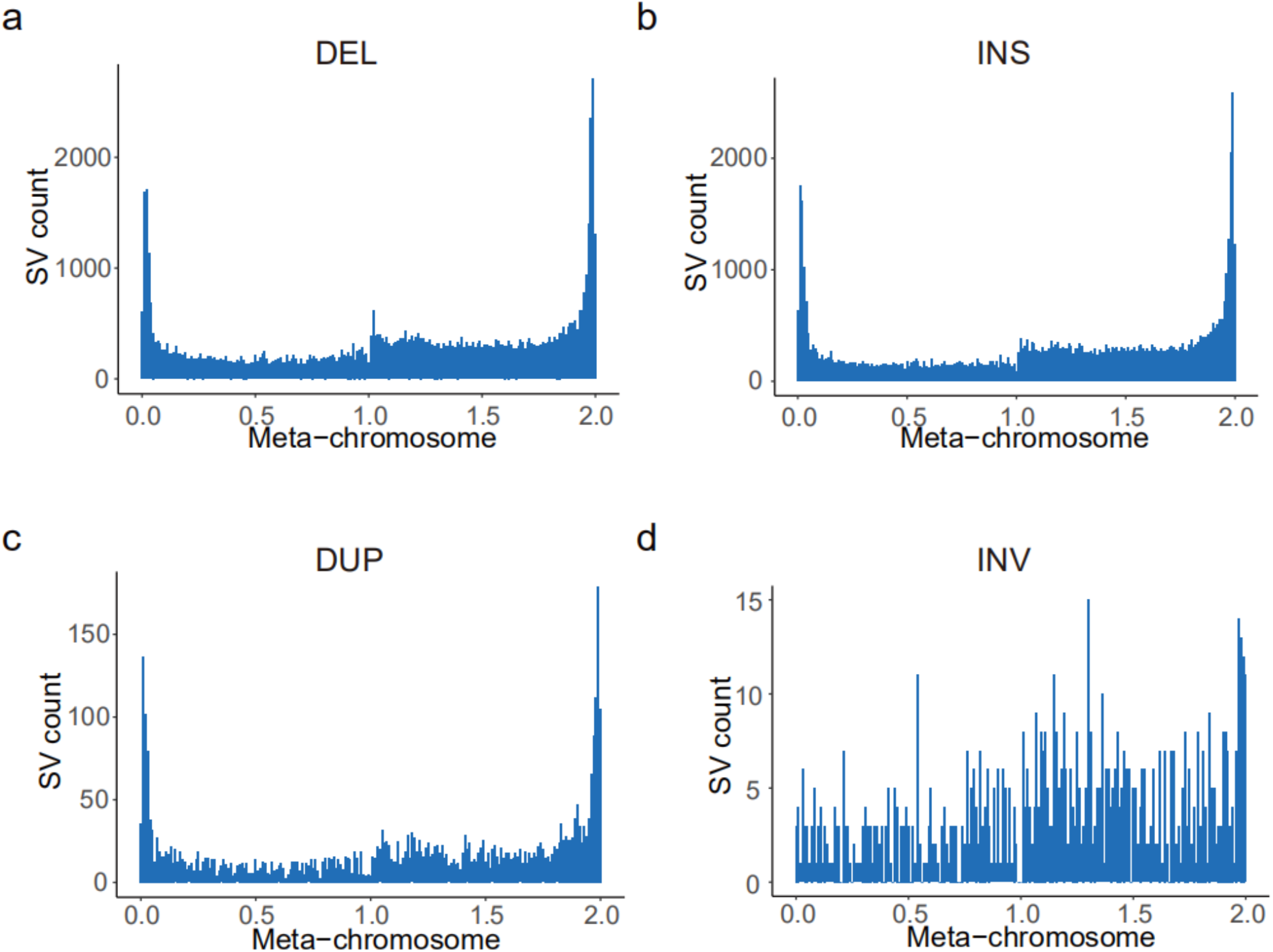
SV distribution across meta-chromosome. SV number across the normalized meta-chromosome for DEL (**a**), INS (**b**), DUP (**c**) and INV (**d**).

**Supplementary Figure 4.**
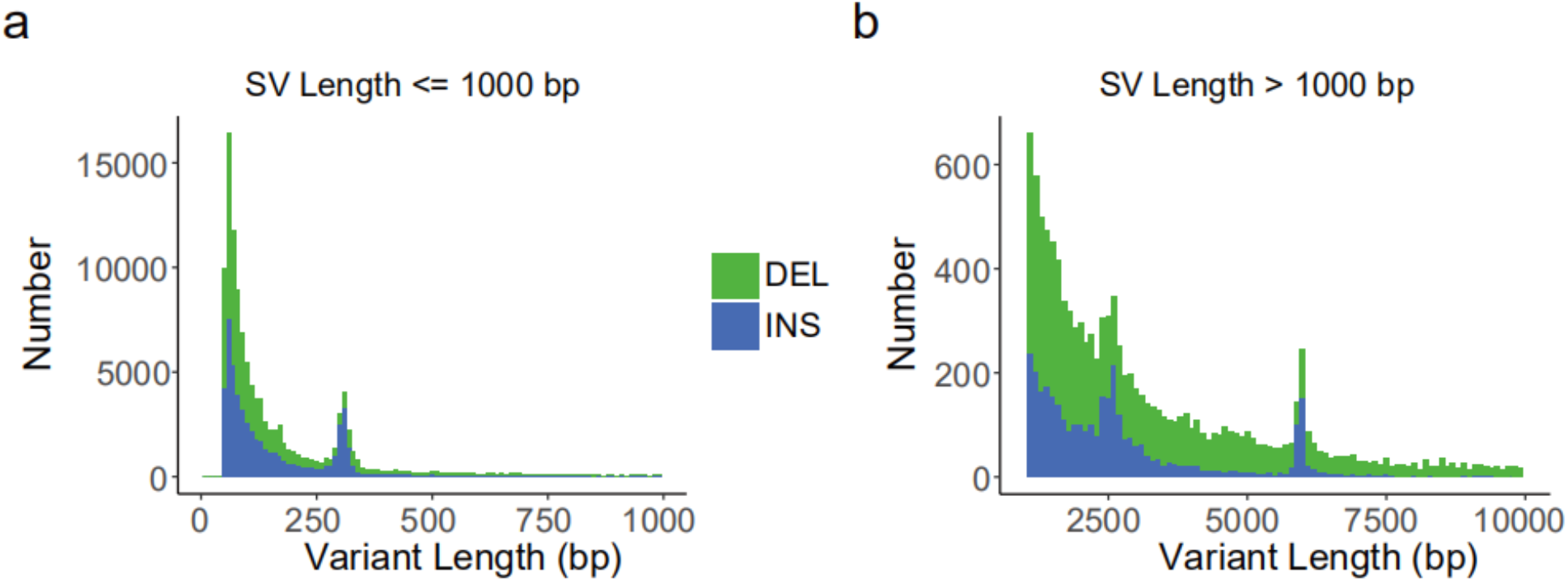
Length distribution for DELs and INSs. SV distribution of DELs and INSs for range of 50 bp to 1 kp (**a**) and range of 1 kb to 10 kb (**b**). Two noticeable peaks were observed at sizes around 300 bp and 6 kb.

**Supplementary Figure 5.**
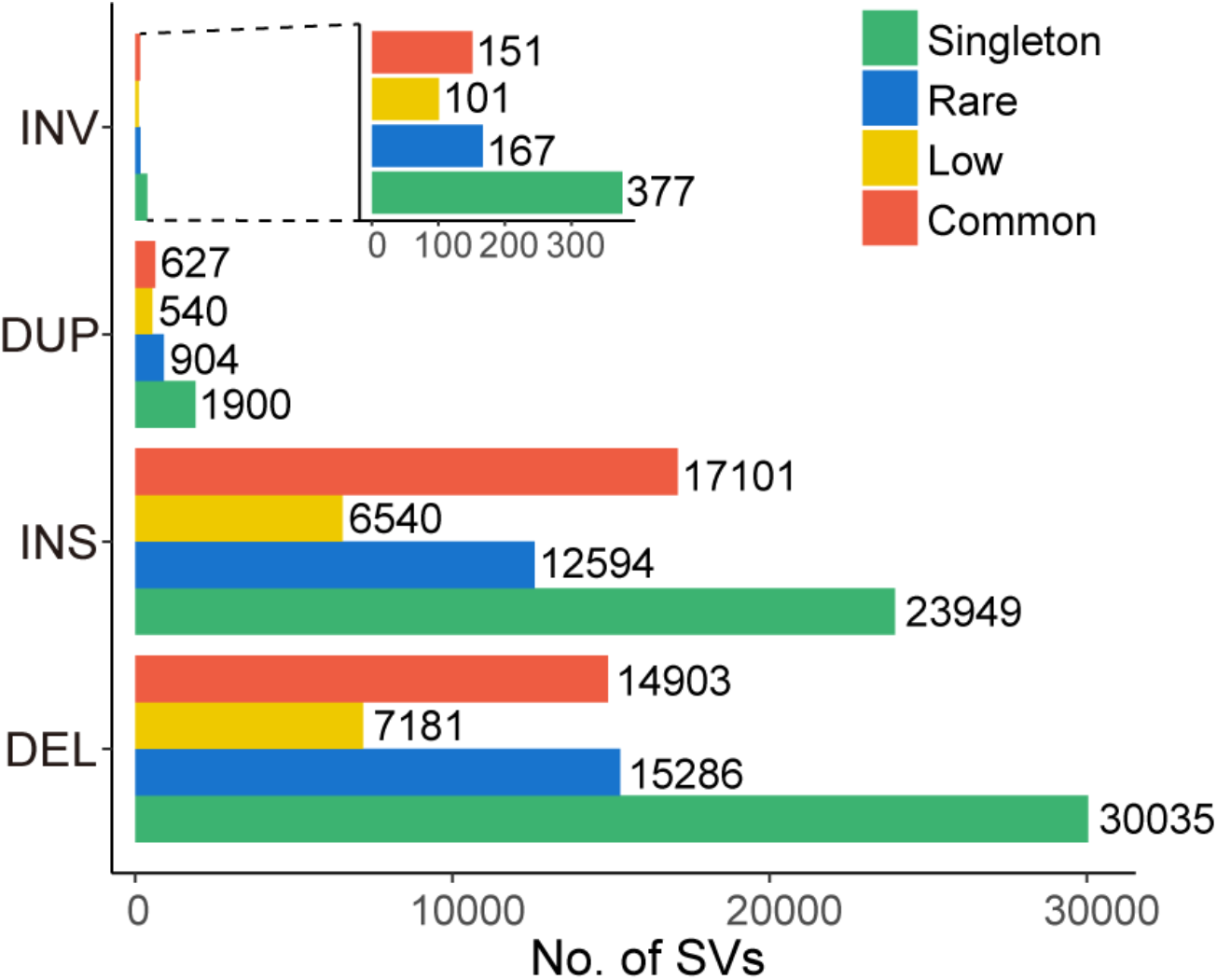
SV numbers of different types in each SV category. SV number for different types of each SV category, the number on the bar graph indicates the actual number of SVs.

**Supplementary Figure 6.**
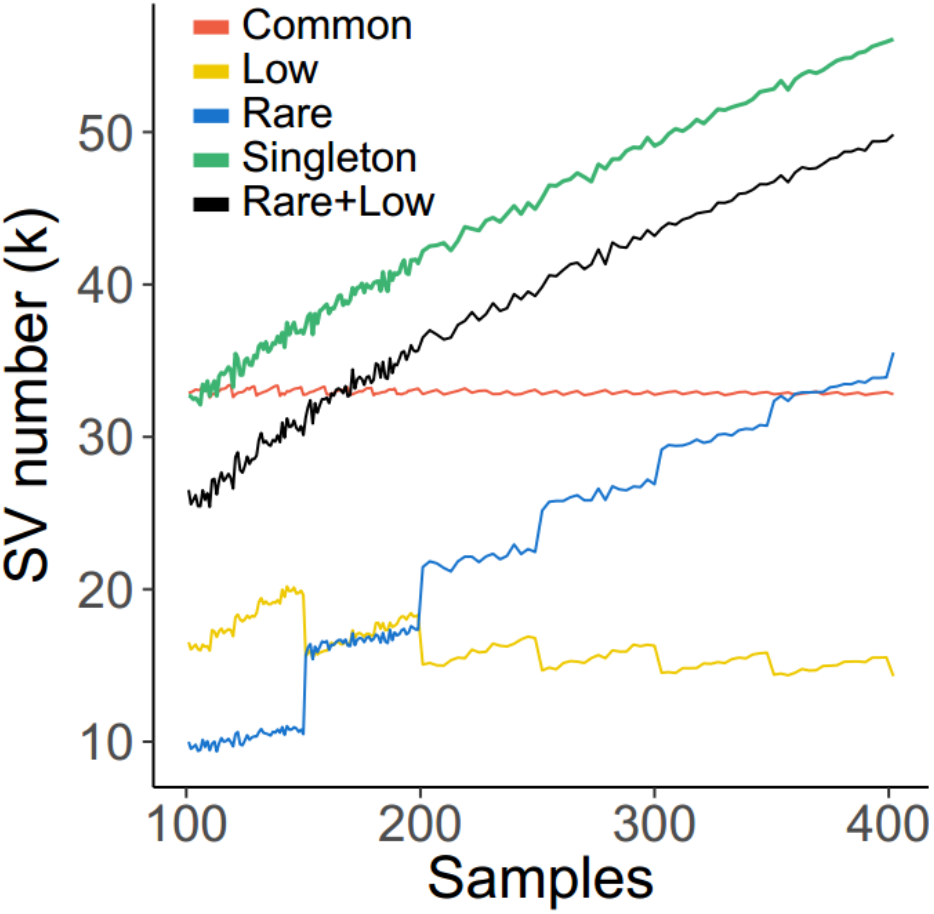
The total number of discovered SVs as a function of number of samples used for detection.

**Supplementary Figure 7.**
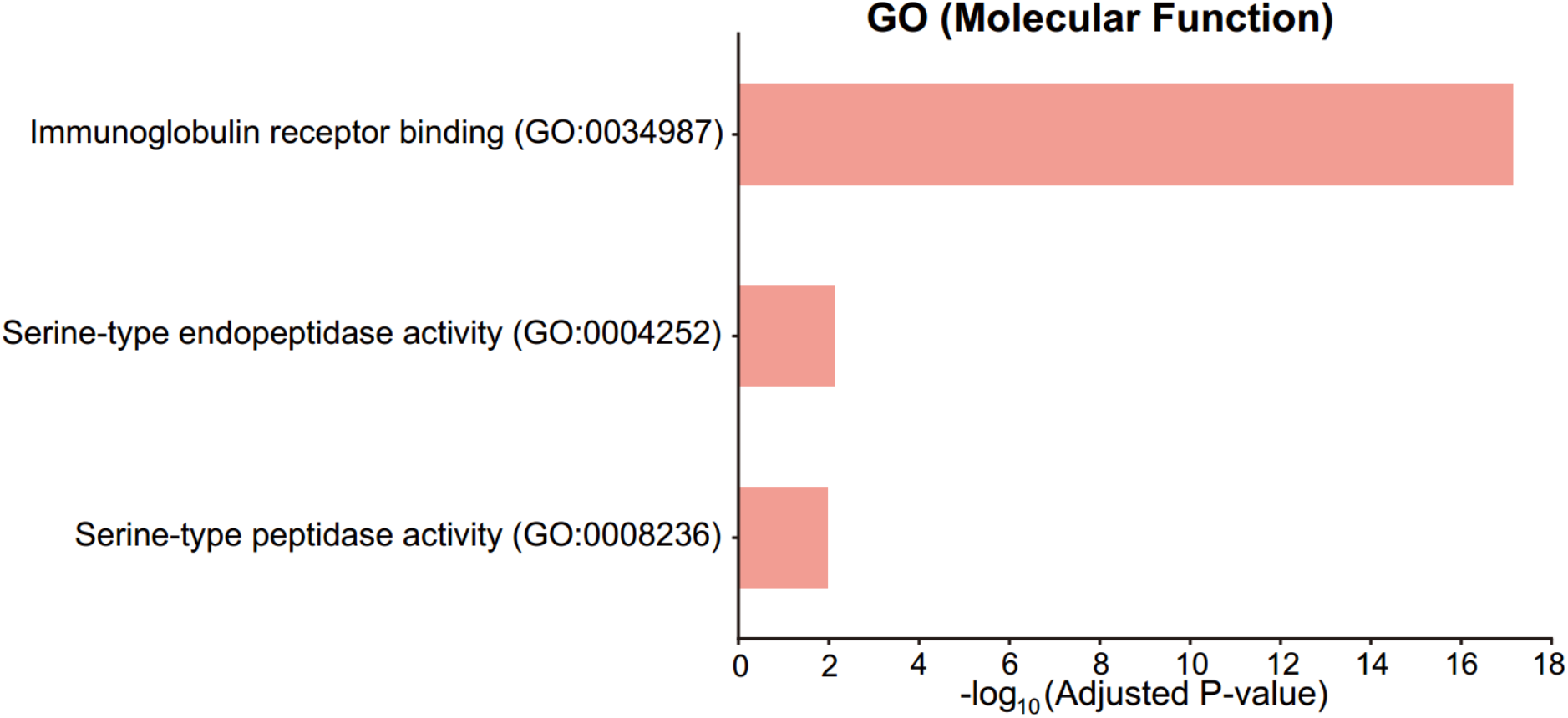
GO enrichment analysis for pLoF SVs associated genes.

**Supplementary Figure 8.**
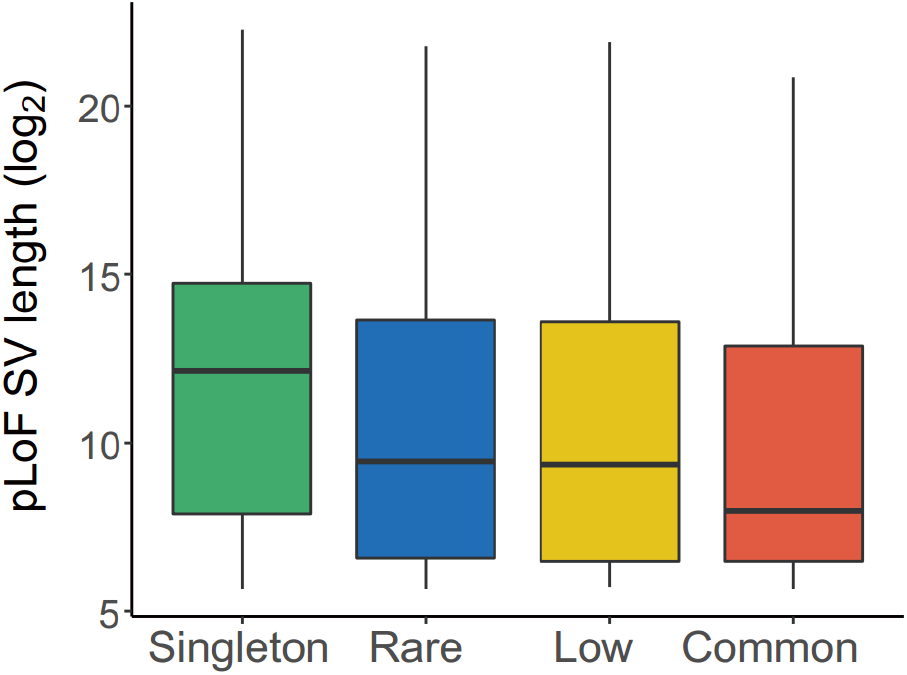
Length distributions of pLoF SVs with different categories.

**Supplementary Table 1. Summary of sample information and sequencing statistics**

**Supplementary Table 2. Summary of clinical traits**

**Supplementary Table 3. SV statistics of different filtering processes for each sample**

**Supplementary Table 4. Accumulative lengths of different type SVs for each sample**

**Supplementary Table 5. Annotation of repeat sequences in SVs**

**Supplementary Table 6. Summary of PCR validation results of singletons**

**Supplementary Table 7. SVs and associated genes in GWAS, OMIM and COSMIC datasets**

**Supplementary Table 8. Association for SVs and clinical phenotypes**

**Supplementary Table 9. Information of SVs with significant *F_st_* (> 0.066) between Southern and Northern Chinese**

